# A versatile, marker-free platform for life cycle-wide imaging in *Plasmodium falciparum* via integration of an exogenous gene cassette into a conserved intergenic locus

**DOI:** 10.64898/2026.01.21.700814

**Authors:** Takashi Sekine, Naoaki Shinzawa, Rie Kubota, Daisuke Kobayashi, Yawara Okubo, Kentaro Itokawa, Haruhiko Isawa, Hisako Amino, Tomoko Ishino

## Abstract

The development of a transgenic *Plasmodium falciparum* line that exhibits robust fluorescence throughout its life cycle in cultured cells and mosquito hosts is valuable for live imaging, tissue localization studies, and quantitative evaluations of parasite infectivity. In this study, we utilized *Plasmodium*-optimized genome editing to integrate an mCherry expression cassette into a specific intergenic locus without gene disruption; thereby successfully minimizing the physiological impact of exogenous gene introduction. The resulting transgenic parasite line, NF54-mCh, exhibited intense fluorescence across all developmental stages; including intraerythrocytic asexual and sexual blood stages; mosquito ookinete, oocyst, and sporozoite stages; and liver stages. Cultured NF54-mCh exhibited normal intraerythrocytic proliferation, gametocytogenesis, and gametocyte maturation, with efficient transmission to mosquito vectors. The marker brightness in salivary gland sporozoites allowed for visual, non-invasive identification of infected mosquitoes, thus facilitating dissection. Sporozoites dissected from salivary glands were highly infectious to livers in humanized mice; and allowed completion of the full life cycle, as evidenced by the appearance of ring-stage parasites in inoculated human erythrocytes. NF54-mCh serves as a parental line for performing additional genetic modifications, because the CRISPR/Cas9-based genome editing method is free of introduced drug resistance markers. The broader applicability of this strategy was validated by generating fluorescence reporter lines for use in the rodent malaria parasite model systems, *Plasmodium berghei* and *Plasmodium yoelii*. In summary, NF54-mCh represents a unique, versatile platform that will accelerate fundamental research and support the future development of innovative malaria control strategies, including studies on new vaccines and drug efficacy.

**IMPORTANCE:** A detailed understanding of the molecular and cellular biology of the human malaria parasite *Plasmodium falciparum*, particularly during infection processes, is essential for efforts in malaria control and elimination. In this study, we generated a marker-free transgenic *P. falciparum* line which exhibits constitutive, robust fluorescence throughout its developmental stages in erythrocytes, mosquitoes, and hepatocytes, without impairing parasite growth or fitness. The fluorescence marker gene was inserted into an intergenic locus and thus no genes were disrupted. The *pfs47* gene is intact, which allows the parasites to evade the mosquito innate system; therefore, the parasite retains the ability for *pfs47*-dependent infection of refractory *Anopheles* strains. The transgenic parasites do not contain drug resistance markers, and thus there are no limitations for further genetic manipulation. This transgenic parasite line provides a powerful tool for the study of fundamental parasite infection mechanisms and for the molecular analysis of host-parasite interactions.

## INTRODUCTION

Malaria is caused by bloodstream infection with the protozoan *Plasmodium* parasite, and globally is a major infectious disease with 263 million cases and 597,000 deaths in 2023, according to WHO reports (1). The emergence of drug resistance of blood stage *Plasmodium falciparum* to first-line antimalarials has become a significant public health concern for malaria control. The first malaria vaccine, RTS,S, which targets sporozoites, was approved by the WHO in 2021 (2) and presently is provided to children in African countries. However, its efficacy to prevent malaria infection appears to be limited and further improvements and vaccines are awaited. To develop new interventions to eradicate malaria, it is necessary to understand the cellular and molecular biology of *P. falciparum*; especially during events such as parasite invasion of target cells, including hepatocytes and erythrocytes in humans, and epithelial cells and salivary glands in mosquitoes. The molecular mechanisms of *Plasmodium* have long been studied using genetically modified *P. falciparum* or rodent malaria parasites. However, human and rodent parasites show differing features suggestive of distinct evolutionary histories; such as dissimilar kinetics of parasite proliferation and differentiation during the intraerythrocytic and liver stages. For the development of new drug therapies or vaccines, it is ultimately necessary to conduct experiments with *P. falciparum* rather than rodent malaria parasite model systems. To monitor parasite development and to evaluate parasite infection, a *P. falciparum* parasite line which expresses a cytosolic fluorescent protein throughout its life cycle and possesses a wild type phenotype would be a powerful research tool.

For the improvement of infection-blocking vaccines, recent studies in the field of *P. falciparum* research have focused on mosquito stage analysis using blood feeding experiments and *in vitro* liver stage analysis using cultured hepatocytes or liver organoid culture (3–8). Research is now being conducted utilizing immunodeficient humanized mice with transplanted human hepatocytes for *in vivo* analysis (9–13); thereby expanding the methods available for analyzing the transmission mechanisms of the human-specific *P. falciparum* (14–20). For localization and quantitative analyses in these liver models, it is advantageous to use parasites that constitutively express fluorescent proteins (21). In past conventional methods, marker genes were integrated within select target genes which were dispensable for parasite development. For example, all *P. falciparum* lines that constitutively express fluorescent proteins and have been used for liver stage analysis were constructed with the disruption of the *pfs47* gene (22–26).

However, because *pfs47* plays a role in evasion mechanisms from the mosquito immune system (27), it may not be a suitable platform for some investigations on mosquito-parasite interactions. In a previous report, it was shown that the introduction of an exogenous gene into an intergenic locus on chromosome 6 of *Plasmodium berghei* can produce a parasite strain that is phenotypically silent and emits green fluorescence (28). Therefore, we decided to insert a fluorescent protein expression cassette into a sufficiently long intergenic locus in the *P. falciparum* NF54 strain to minimize the effects of transgenesis without gene disruption.

We additionally were interested in addressing the issue of the drug resistance cassette, which remains in most reported fluorescence-expressing *P. falciparum* parasites (24). The advent of CRISPR/Cas9 technology has significantly improved the genome editing efficiency and ease of working with *P. falciparum* parasites, which historically required more difficult genetic modification methods than rodent malaria parasite model systems (29–31). While a few marker-free lines have been developed by these highly efficient genome editing techniques, they often exhibit limitations such as reduced infectivity in liver stages or inconsistent fluorescence intensity across life cycle stages or impaired infectivity. The generation of fluorescent *P. falciparum* parasites that are both marker-free and highly fluorescent at all stages is expected to facilitate the evaluation of the effects of gene knockouts and other modifications, thereby accelerating future elucidations of infection mechanisms.

To address the above concerns, this study describes the generation of a *Plasmodium falciparum* parasite line, designated NF54-mCh, that constitutive expresses an mCherry fluorescent protein throughout its life cycle. Unlike traditional lines targeting the *pfs47* gene, by integrating an mCherry cassette into a conserved intergenic locus on chromosome 10 using an improved CRISPR/Cas9 system, our strategy preserved the immune evasion machinery in the mosquito vector. The resulting parasites exhibited intense fluorescence across the life cycle; and the remarkable brightness in salivary glands facilitated the non-invasive identification of infected mosquitoes. This marker-free strain can serve as a parental line for further genetic modifications using drug-selectable markers. We also validated the broader utility of this intergenic locus by successfully generating a reporter-expressing parasite line in the rodent malaria parasites *Plasmodium berghei* and *Plasmodium yoelii*. Thus, this study provides a robust, versatile genetic platform that not only enhances our ability to perform high-resolution fluorescence imaging but also establishes a transformative tool for the future development of innovative malaria control strategies, including studies on new vaccines and drugs.

## RESULTS

### Generation of a *P. falciparum* line which constitutively expresses mCherry

To avoid gene disruption and minimize the impact of inserting an mCherry expressing cassette, we selected a region on chromosome 10 between the gene loci PF3D7_1022300 and PF3D7_1022400. These genes are 4 kb apart with convergent gene orientation, and both are reported as dispensable genes in asexual stages (32). The *pfhsp90* promoter was chosen to drive an mCherry expression cassette, in anticipation that following integration into the intergenic locus using the CRISPR/Cas9 system it would express the fluorescence marker in all *P. falciparum* mosquito and human life cycle stages (Figure 1A). Diagnostic genotyping PCR of the obtained transgenic parasites, designated as NF54-mCh, indicated that the mCherry cassette was introduced to the expected location (Figure 1B). In the population obtained after drug selection, the wild-type DNA fragment was barely detectable by PCR, suggesting efficient genome editing. Two parasite clones were obtained by limiting dilution, in which only the mCherry expression cassette was integrated at the targeted intergenic locus, and were named NF54-mCh (B2) and NF54-mCh (D6). Southern blotting analysis confirmed the integration of the mCherry expression cassette exclusively at the expected locus (Figure 1C). Whole-genome sequencing of NF54-mCh (B2) was performed to assess the possibility that the CRISPR/Cas9 editing introduced unexpected mutations which might affect the parasite phenotype. A total of 13 indels and 2 SNPs unique to NF54-mCh were identified (Supplemental Table 1). All indels were located in AT-rich intergenic regions and an intron, and are therefore unlikely to impact gene expression. The two SNPs were found within the repeat region of PF3D7_0322700 but were synonymous mutations. Therefore, whole genome sequencing of NF54-mCh indicated that the insertion of the mCherry expressing cassette did not affect genomic DNA at any loci.

**Fig. 1.**
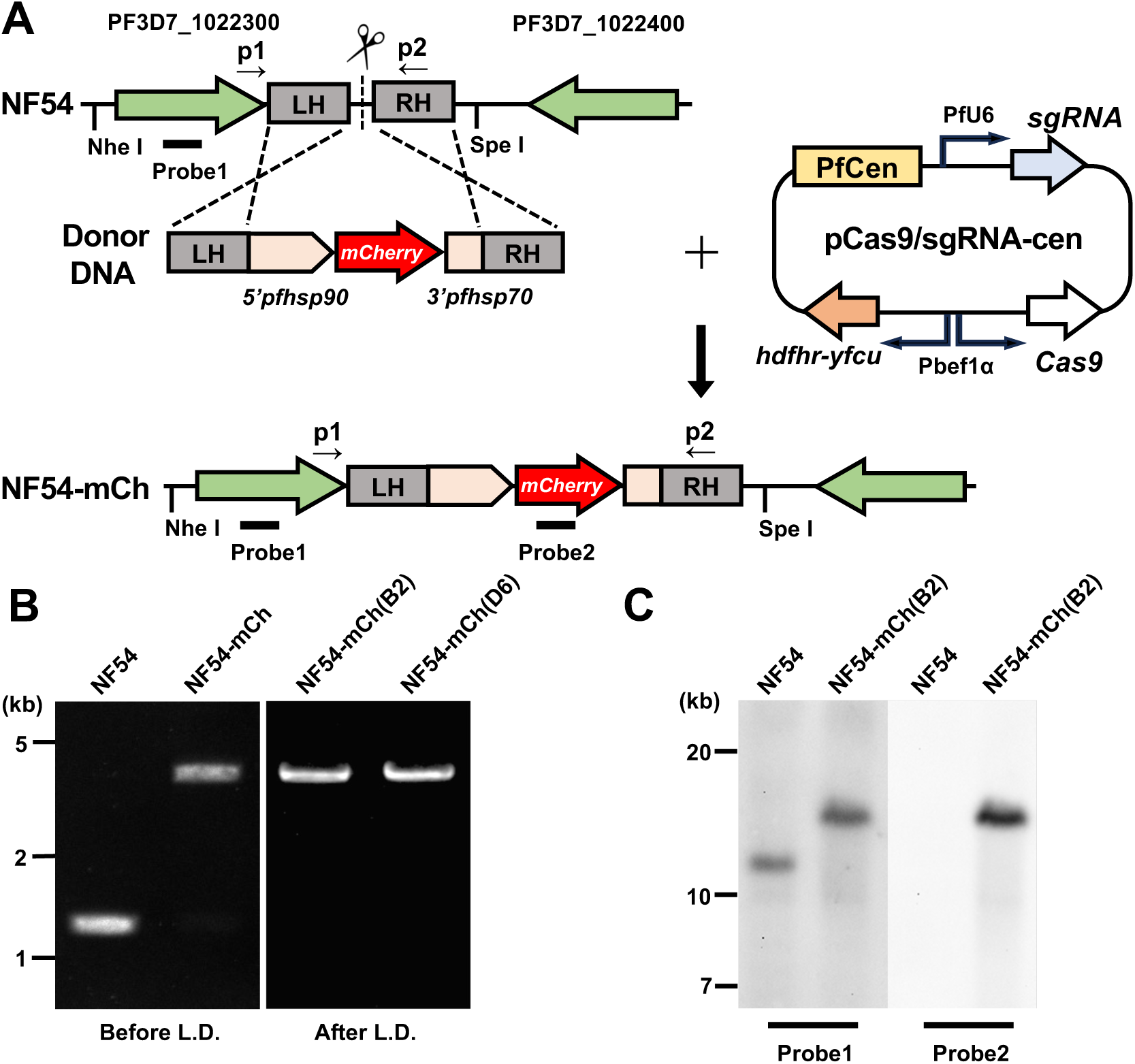
Generation of mCherry expressing *P. falciparum* NF54. (A) A schematic representation of genome editing using the CRISPR/Cas9 system to integrate an mCherry expressing cassette into an intergenic locus of NF54 strain parasites. The donor DNA fragment (Donor) consisted of two homologous regions (LH and RH) and an mCherry coding region with a *pfhsp90* promoter and a *pfhsp70* 3’UTR region. Plasmid DNA (pCas9/sgRNA) contained a Pf centromere sequence (PfCen) which expresses Cas9 and hDHFR-yFCU under the *pbef1a* promoter together with guide RNA (sgRNA) under a *pfU6* promoter. The DNA integrated locus after transfection of these DNA fragments and plasmid into NF54 parasites is demonstrated below (NF54-mCh). The PCR primers p1 and p2 shown in Fig. 1B correspond to pfcheck-F and pfcheck-R. Restriction enzyme sites NheI and SpeI, and the probe used for genomic Southern blotting (Fig. 1C) are also indicated. LH, left homologous region; RH, right homologous region. (B) The genotyping PCR of transfected parasites. DNA fragments were amplified using p1 and P2 primers (indicated in A) with genomic DNA of transfected parasites and two clones obtained from limiting dilution. Longer fragments demonstrated that the mCherry expression cassette was correctly integrated at the indicated locus. The DNA size marker was indicated on the left. (C) Genomic Southern blot hybridization analysis of NF54-mCh parasites. Genomic DNAs of NF54 and cloned NF54-mCh parasites were digested with NheI and SpeI, electrophoretically separated, and hybridized with either probe1 or probe2 indicated in A. The signals detected at approximately 18 kbp correspond to the genomic DNA fragment size after genome editing at the precise locus. A DNA size marker is indicated on the left.

### Fluorescence in NF54-mCh asexual stages

The proliferation rate of cultured NF54-mCh blood stage parasites was examined; and the growth of the two clones were comparable to the parental line NF54 (Figure 2A). mCherry fluorescence was observed in the cytosol at each intraerythrocytic stage of NF54-mCh parasites by live-imaging using fluorescence microscopy (Figure 2B). Thus NF54-mCh parasites express a strong mCherry signal throughout intraerythrocytic parasite development without affecting the growth rate.

**Fig. 2.**
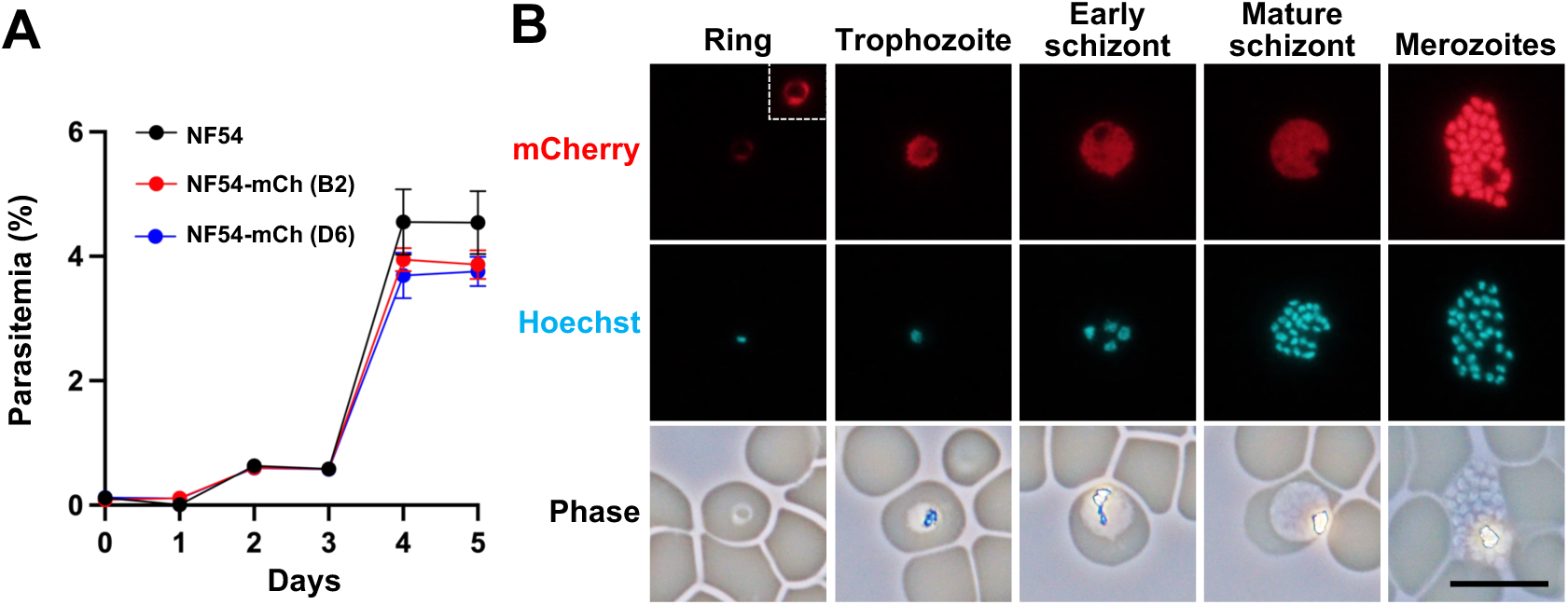
Asexual development of NF54-mCh parasites. (A) The comparison of parasite growth *in vitro* between NF54 parental and NF54-mCh parasites. Parasitemias of two clones of NF54-mCh parasites (red and blue) were comparable to those of the parental NF54 parasites (black). Dots indicate the mean values with standard deviations as error bars from three independent experiments. There were no significant differences in parasitemias on each day among the parasite lines using the unpaired *t*-test. (B) mCherry expression during asexual development in NF54-mCh. Thin smears of NF54-mCh infected erythrocytes were made without fixation, and mCherry fluorescence (red, top panels) was captured using a fluorescence microscope. All mCherry images were captured with the same exposure time, except for an image in the insert, which was taken with a longer exposure time to visualize its signal at the ring stage (upper-right in “Ring”). Nuclei were stained with Hoechst 33342 (cyan, middle panels). Phase images are shown below. The bar indicates 10 µm.

### Normal development of gametocytes was observed in the NF54-mCh parasite

To examine the effect of mCherry expression on sexual stage development, gametocytes were induced from *in vitro* asexual stage cultures by adding human serum without supplying new erythrocytes for 17 days. Gametocytemias were measured from day 10 post induction by Giemsa staining of blood smears on glass slides; and were comparable between the NF54-mCh clones and the parental NF54 line (Figure 3A). An immunofluorescent assay (IFA) using anti-α-tubulin and anti-PfG377 marker antibodies was performed to calculate the proportion of female and male gametocytes. Specifically, male gametocytes showed stronger α-tubulin signals, while female gametocytes showed stronger PfG377 signals in their cytosol (33); and thusly the female/male ratios in the NF54-mCh clones were observed to be comparable to the parental line (Figure 3B). mCherry fluorescence was clearly detected throughout gametocyte maturation (Figure 3C).

**Fig. 3.**
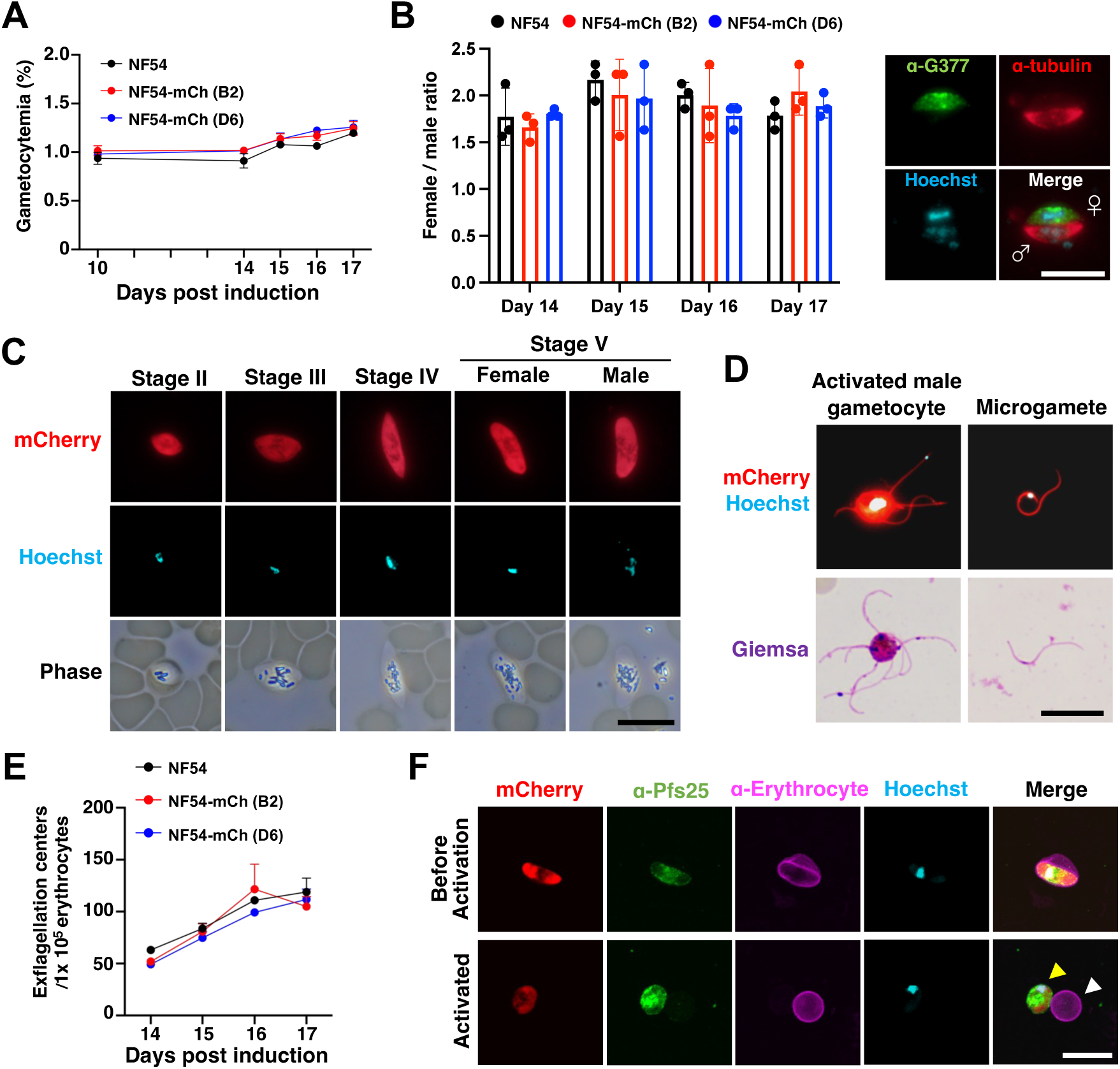
Development and maturation of NF54-mCh gametocytes. (A) The gametocytemias of NF54-mCh parasites after induction during *in vitro* culture. Bars show the means of three independent gametocyte cultures, and error bars indicate standard errors. Distributions were compared using the unpaired *t*-test (not significant). (B) (Left) The ratio of male and female gametocytes within NF54-mCh cultures. Bars show the means of three independent gametocyte cultures, and error bars indicate standard errors. Distributions were compared using the unpaired *t*-test and were not significant. (Right) Male and female gametocytes were detected with rabbit anti-PfG377 antiserum (green) and mouse anti-α-tubulin monoclonal antibody (red), respectively. Nuclei were stained with Hoechst 33342 (cyan). The bar indicates 10 µm. (C) Representative images during NF54-mCh gametocyte maturation. Thin smears of NF54-mCh infected erythrocytes were made without fixation, and mCherry fluorescence (red, top panels) was captured using a fluorescence microscope. Nuclei were stained with Hoechst 33342 (cyan, middle panels) and phase images (lower panels) are shown. The indicated gametocyte stages were determined by their morphologies. The bar indicates 10 µm. All images were captured with the same exposure time. (D) Representative images of an activated male gametocyte and a microgamete of NF54-mCh. Cultured parasites were incubated with human serum for 15 minutes before making thin smears. mCherry fluorescence (red) together with nuclei stained with Hoechst 33342 (cyan) are shown in the upper panels. Giemsa staining of activated parasites are shown in the lower panels. The bar indicates 10 µm. (E) The number of exflagellation centers of the NF54-mCh after *in vitro* gametocyte activation. The mean number of exflagellation centers per 10^5^ erythrocytes from three independent experiments using NF54 (black) and two NF54-mCh clones (red and blue) are shown as symbols with error bars indicating standard errors. The day of activation after gametocyte culture is indicated below. No significant difference was observed in the number of activated male gametocytes throughout the days examined by the unpaired *t*-test. (F) The evaluation of female gametocyte egress from erythrocytes. Cultured NF54-mCh parasites containing mature gametocytes were incubated with or without human serum for 15 minutes before fixation with 4% paraformaldehyde. Female gametocytes and erythrocyte membranes were detected by mouse anti-Pfs25 monoclonal antibodies (green) and rabbit anti-human erythrocyte polyclonal antibodies (magenta), respectively. Nuclei were stained with Hoechst 33342 (cyan). Merged images are shown in the right panels. Upon activation, a female gametocyte egresses from an erythrocyte (yellow arrowhead, lower panels). The white arrowhead indicates uninfected red blood cells. The bar indicates 10 µm.

To examine the transmission ability of NF54-mCh parasites to mosquitoes, firstly the number of exflagellation centers were counted as an indicator of the capacity of male gametocytes to differentiate into gametes after being ingested by mosquitoes. Gametogenesis was induced by incubating mature gametocytes in human serum for 15 minutes at room temperature (34), and then the released microgametes expressing mCherry were counted using a microscope (Figure 3D). The numbers of exflagellation centers of NF54-mCh from days 14 to 17 post gametocyte-induction were comparable to those of the NF54 parental line (Figure 3E). Female gametocyte activation was measured by IFA using anti-Pfs25 and anti-human erythrocyte antibodies. As shown in Figure 3F, activated, rounded mCherry expressing macrogametes were observed which were egressed from erythrocytes. These results indicated that NF54-mCh parasites differentiated normally into sexual stage parasites; and strongly expressed cytosolic fluorescence.

### NF54-mCh parasites develop normally in *Anopheles* mosquitoes

For *P. falciparum*, gamete fertilization and ookinete transformation cannot be fully replicated *in vitro*, making it necessary to analyze the development of these stages using the mosquito host (35). This caveat in turn emphasizes that fluorescence-expressing parasites are a particularly useful tool for monitoring infection and localization during parasite invasion and maturation in mosquito vectors. NF54 and NF54-mCh parasite transmission to *Anopheles stephensi* mosquitoes was performed by membrane-feeding using gametocytes at day 17 post-induction. Ookinetes harvested from NF54-mCh infected mosquito midguts at 24 hours post-feeding showed strong cytosolic mCherry signals (Figure 4A). At 8 to 10 days post-feeding, mCherry-positive oocysts were observed on dissected midguts (Figure 4B), and the oocyst numbers were comparable between NF54-mCh and the NF54 parental line (Figure 4C and Supplemental Figure 1). mCherry fluorescence originating from NF54-mCh sporozoites was sufficiently strong to be detected from the outside of salivary glands (Figure 4D and E), as well as from the outside of infected mosquitoes (Figure 4F). Dissection of infected salivary glands at days 18 to 20 post-feeding showed comparable numbers of NF54-mCh versus NF54 wild type sporozoites (Figure 4G). Because sporozoites residing in salivary glands are inoculated into the human skin through mosquito bites, the finding that NF54-mCh and NF54 parental line sporozoites accumulated in salivary glands with similar efficiency demonstrated that the transmission efficacy of NF54-mCh parasites via mosquitoes is likely equivalent to its parental strain.

**Fig. 4.**
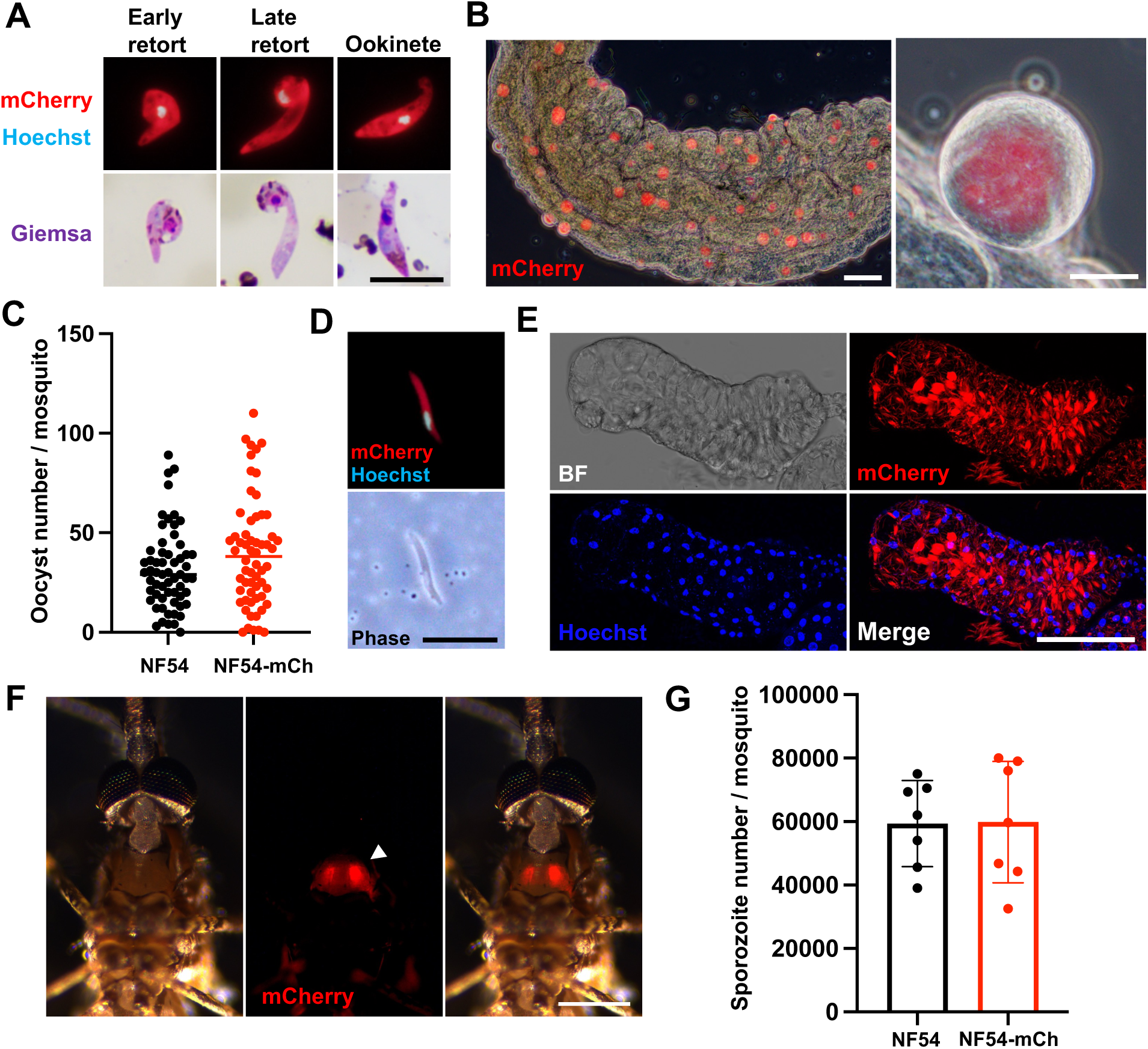
NF54-mCh parasite transmission to mosquitoes. (A) Representative images of mCherry fluorescence during ookinete maturation in mosquito midguts. A homogenate of midguts collected from NF54-mCh infected mosquitoes at day 1 post-feeding was smeared on a glass slide to detect mCherry (red), with nuclei stained with Hoechst 33342 (cyan). Giemsa staining ookinetes are shown below. The bar indicates 10 µm. (B) A representative image of mCherry expression in oocysts on the midgut of an NF54-mCh infected *A. stephensi* mosquito. Midguts were dissected at days 8 to 10 post-feeding and observed using a fluorescence microscope with a phase image. The bar in the left panel indicates 100 µm, and the right indicates 20 µm. (C) The comparison of oocyst number on mosquito midguts between the NF54 parental line and NF54-mCh. Midguts were collected at days 8 to 10 post-feeding and visualized by staining with mercurochrome. Each dot represents the number of oocysts from an individual midgut, with horizontal lines indicating the median. The Mann-Whitney test was used to compare distributions (N = 60). (D) Representative images of an mCherry expressing sporozoite. Nuclei were stained with Hoechst 33342 (cyan). The bar indicates 10 µm. (E) Micrograph of a confocal section of a sporozoite-residing salivary gland. A salivary gland from an NF54-mCh infected mosquito was collected at days 18 to 20 post-feeding and stained with Hoechst 33342 to visualize gland nuclei (blue). Fluorescence images are shown as max-intensity projections of confocal stacks. The bar indicates 100 µm. (F) External view of a live *A. stephensi* mosquito showing salivary gland fluorescence. The picture was captured at days 16 post-feeding. The arrowhead indicates mCherry signals in the salivary glands. A scale bar indicates 500 µm. (G) The number of sporozoites collected from mosquito salivary glands at days 18 to 20 post-feeding. Seventy pairs of salivary glands were collected from NF54 or NF54-mCh infected mosquitoes to count sporozoites that successfully invaded salivary glands. Each dot represents the number of pooled sporozoites from ten mosquitos. Bars show the mean numbers from biological septuplicate with error bars indicating standard deviations. The Mann-Whitney test was used to compare distributions (not significant).

### NF54-mCh sporozoites successfully infect human hepatocytes to release infective hepatic merozoites in a humanized mouse model

To assess the ability of hepatocyte infection, NF54-mCh sporozoites dissected from mosquito salivary glands were inoculated onto the hepatocyte-derived immortalized cell line, HC-04, which supports *P. falciparum* liver stage parasite development (36). NF54-mCh sporozoites were capable of infecting HC-04, and showed nuclei division within parasitophorous vacuoles (Figure 5A). Strong cytosolic expression of mCherry was observed in the NF54-mCh liver stage parasites. The infectivity of NF54-mCh hepatic merozoites to erythrocytes was then examined; and thereby conceptual completion of the parasite life cycle. While the production of infectious hepatic merozoites *in vitro* remains challenging and inconsistent, an *in vivo* infection system using human liver chimeric mice, in which hepatocytes are replaced by human hepatocytes, can be applied for the evaluation of the infectivity of hepatic merozoites (6). To account for individual variability among mice, a total of 1.0×10^5^ sporozoites, consisting of a mixture of NF54-mCh and NF54 sporozoites at a ratio of 10:7, were intravenously inoculated into a single humanized mouse. At days 6 and 8 post sporozoite inoculation, human erythrocytes were intravenously inoculated into the infected mouse, and then blood was collected 4 h after the second erythrocyte inoculation to serve for *in vitro* parasite culture (Figure 5B). One day after the initiation of the *in vitro* culture, both mCherry-positive and negative ring forms were detected within blood smears (Figure 5C). On day 4 after establishment of the *in vitro* culture, parasites were counted and the ratio of NF54-mCh parasites to NF54 parasites was 10:7 (116 NF54-mCherry and 84 NF54 parasites out of 200 total parasites counted). Since the proportion of parasites that emerged in the blood was comparable to the ratio of the inoculated sporozoites, it was concluded that there is no significant difference between NF54-mCh and the NF54 parental line with respect to sporozoite infectivity or the efficiency of intrahepatocytic development.

**Fig. 5.**
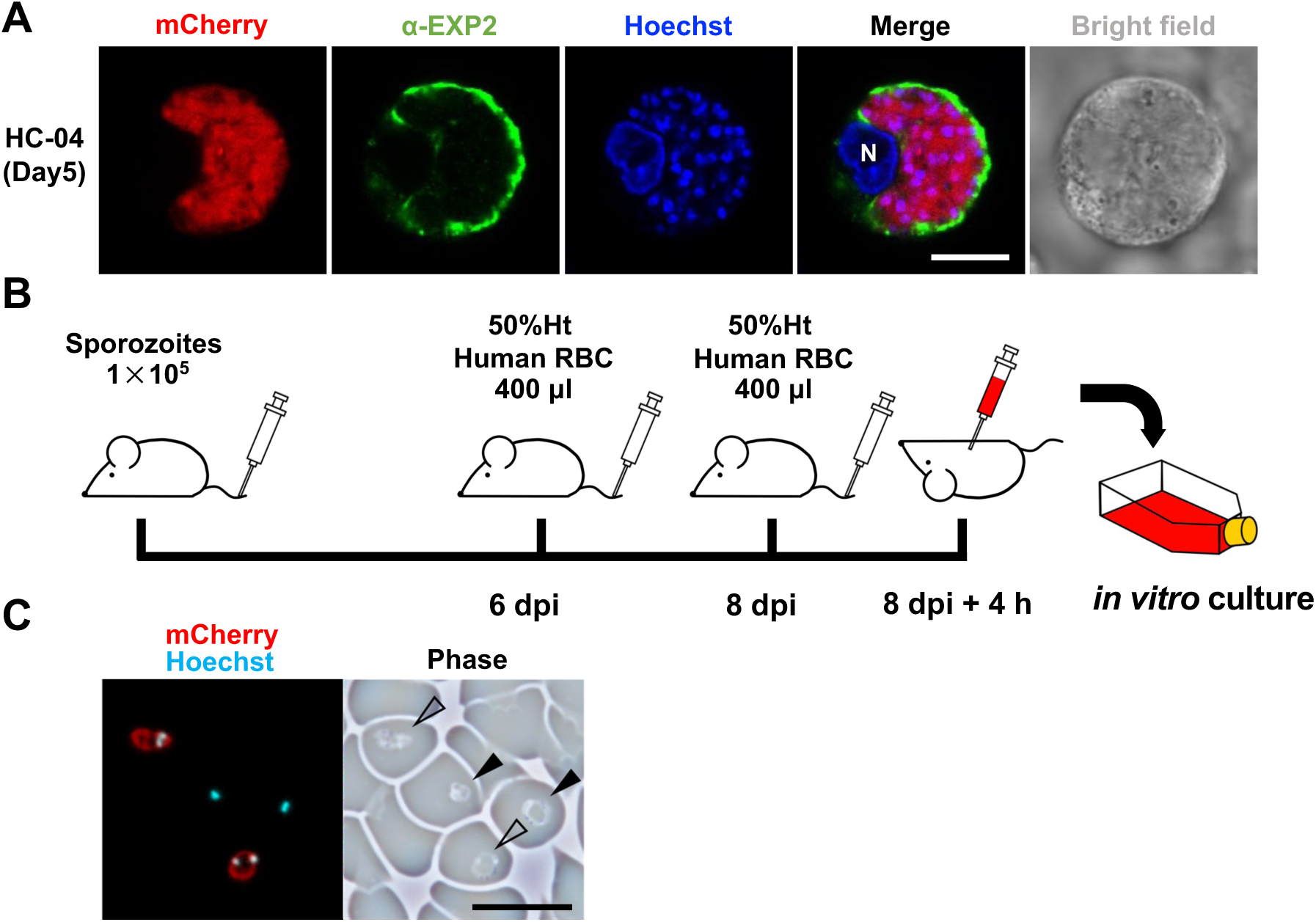
NF54-mCh sporozoite infection of hepatocytes. (A) NF54-mCh parasite development inside hepatocytes. HC-04 cells were fixed with 4% paraformaldehyde 5 days after NF54-mCh sporozoite inoculation. Cells were labeled with rabbit anti-EXP2 antibodies (green), together with Hoechst 33342 (blue). Micrographs of a confocal section of the liver stage parasite are shown. N indicates the host cell nucleus. The bar indicates 10 µm. (B) Workflow to evaluate Pf sporozoite infectivity to a human liver chimeric mouse. Sporozoites collected from salivary glands were intravenously injected into a human liver chimeric mouse, followed by human erythrocytes inoculation at days 6 and 8 post-inoculation. Blood was collected 4 h after the second erythrocyte inoculation, which was subjected to *in vitro* culture in complete medium. (C) Reappearance of blood-stage parasites through the *in vivo* liver stage. Ring stage parasites were detected on the next day from the start of *in vitro* culture indicated at B. Closed arrowheads and open arrowheads indicate NF54 and NF54-mCh parasites, distinguished by mCherry expression (red). Nuclei were stained with Hoechst 33342 (cyan). The bar indicates 10 µm.

### The intergenic locus shows synteny with rodent malaria parasite genomes and can be used for targeted integration of expression cassettes

The potential utility of the intergenic region used for the integration of mCherry in *P. falciparum* was explored in other malaria parasite species as a locus for the integration of exogenous DNA. Specifically, we aimed to insert a reporter protein expressing cassette into the syntenic intergenic region between PBANKA_0506500 and PBANKA_0506600 on chromosome 5 of the rodent malaria parasite, *Plasmodium berghei* ANKA (PbANKA) (Figure 6A). We attempted to express two reporter proteins, mGreenLantern (mGL), a brighter variant of GFP (37), and Akaluc, a red-shifted variant of firefly luciferase (38), fused by a self-cleaving peptide, T2A, and controlled by the universal promoter *pbhsp70*. Genotyping PCR and Western blotting analysis confirmed the correct integration into the syntenic intergenic region; and therefore, we named this transgenic parasite Pb-mGL-Aka (Figure 6B and 6C).

**Fig. 6.**
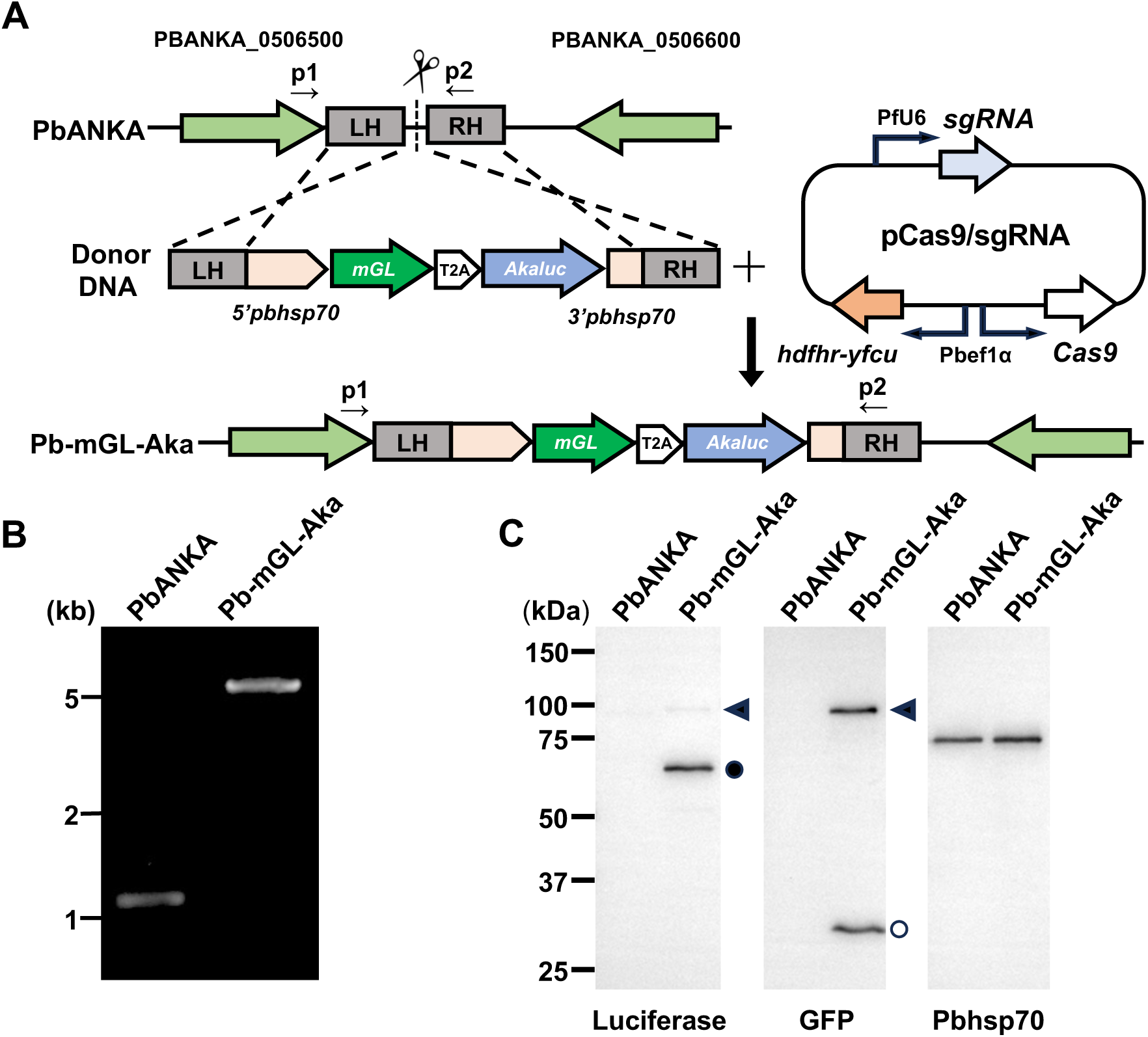
Generation of mGreenLantern-Akaluc expressing *P. berghei*. (A) A schematic representation of CRISPR/Cas9 system genome editing to integrate an mGreenLantern-Akaluc expression cassette into an intergenic locus. The donor DNA fragment consisted of two homologous regions (LH and RH) and an mGreenLantern-Akaluc coding region with a *pbhsp70* promoter and *pbhsp70* 3’UTR. Plasmid DNA (pCas9/sgRNA) which expresses Cas9 and hDHFR-yFCU under a *pbef1a* promoter together with guide RNA (sgRNA) under a *pfU6* promoter. The DNA integrated locus after transfection of these DNA fragments and plasmid into PbANKA parasites is demonstrated below (Pb-mGL-Aka). The PCR primers p1 and p2 shown in Fig. 6B correspond to pbcheck-F and pbcheck-R. LH, left homologous region; RH, right homologous region. (B) The genotyping PCR of transfected parasites. DNA fragments were amplified using p1 and p2 primers (indicated in A) with genomic DNA of transfected parasites and a clone obtained following limiting dilution. Longer fragments demonstrated that the mGreenLantern-Akaluc expression cassette was correctly integrated at the indicated locus. The DNA size marker is indicated on the left. (C) Western blot analysis of Pb-mGL-Aka. Schizonts of PbANKA and Pb-mGL-Aka parasites were collected for the analysis. Samples were adjusted to contain 1×10^6^ parasites and extracts were electrophoretically separated, transferred to membranes, and subjected to Western blot analysis. Antibodies against luciferase and GFP were used to detect the respective proteins. Pbhsp70 antibody was employed as a loading control. Using the anti-luciferase antibody, two bands at approximately 90 kDa and 60 kDa were detected in Pb-mGL-Aka, corresponding to the mGL-Akaluc fusion protein and the cleaved Akaluc protein, respectively. Similarly, probing with the anti-GFP antibody revealed bands at approximately 90 kDa and 30 kDa, representing the fusion protein and cleaved mGL, respectively. Arrowhead, closed circle, and an open circle represent the 90 kDa, 60 kDa, and 30 kDa bands, respectively.

In the Pb-mGL-Aka parasites, green fluorescence of mGL was clearly observed throughout the blood stages as well as in gametocytes (Figure 7A). Fluorescence was also observed in the mosquito stages; that is, ookinetes, oocysts, and sporozoites (Figure 7B-E). The numbers of oocysts and sporozoites were compared with those of the parental line, and no significant differences were observed (Figure 7F, G). To explore whether the liver stage parasites could be measured by luciferase activity from Akaluc, 1.0×10^4^ Pb-mGL-Aka sporozoites were intravenously inoculated into an ICR mouse. Twenty-four hours later, the Akaluc substrate was injected to the mouse and bioluminescence was detected (Figure 7H). Pb-mGL-Aka parasites were able to infect mice and proliferate *in vivo* with comparable efficiency to PbANKA following the intravenous injection of 2000 sporozoites into C57BL/6 mice. (Figure 7I). These results demonstrate that the insertion of an mGreenLantern-Akaluc expression cassette into the intergenic locus in *P. berghei* chromosome 5 drives expression of bright green fluorescence through the life cycle without interfering with *P. berghei* parasite infection and proliferation. Similarly, introduction of the mGL expression cassette into an intergenic locus of *Plasmodium yoelii* resulted in detectable fluorescence in the blood stages, oocysts, and sporozoites (Supplemental Figure 2). These findings support that this intergenic locus serves as a permissive site for the integration of exogenous genes in *P. falciparum*, *P. berghei*, and *P. yoelii*.

**Fig. 7.**
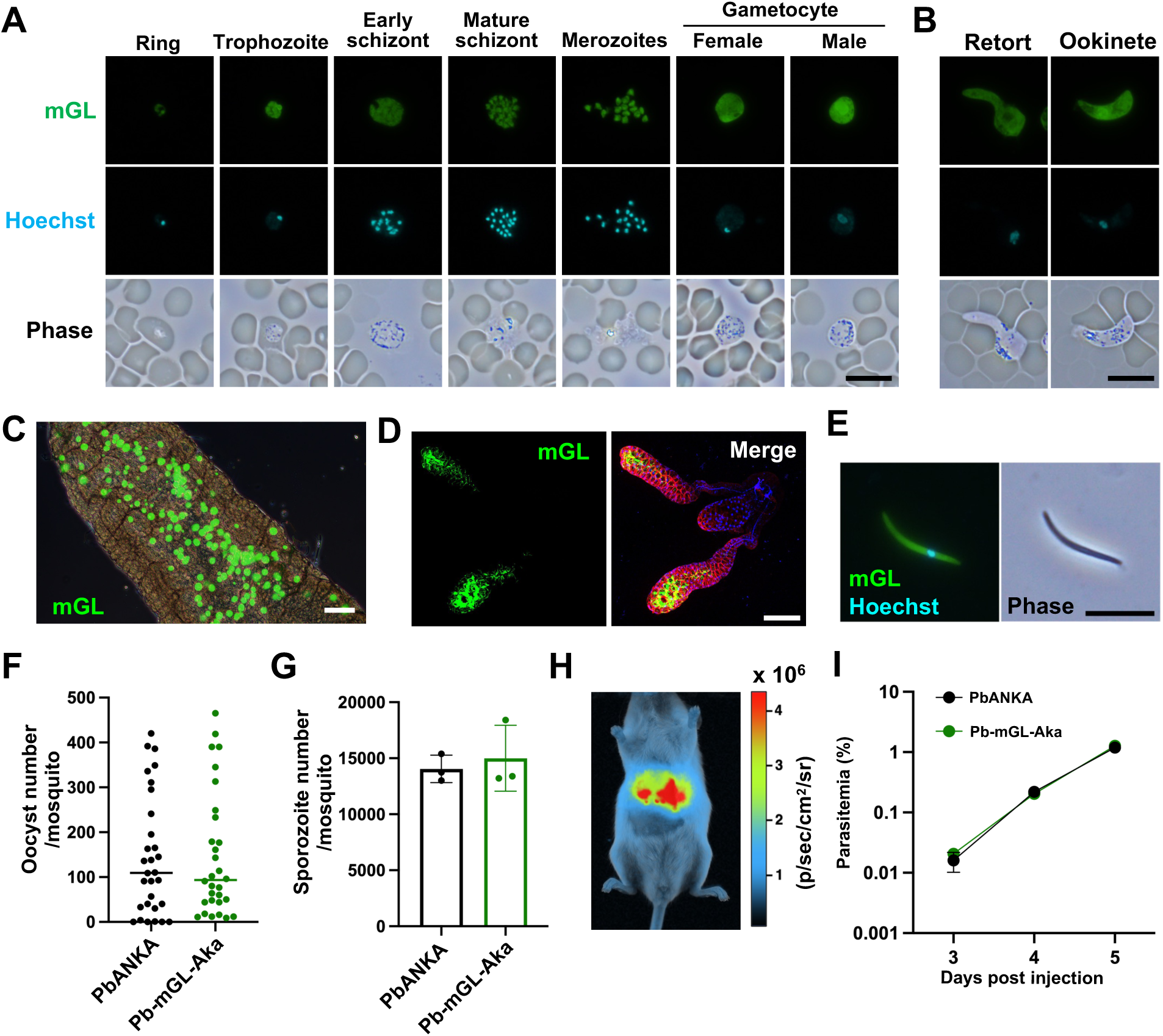
mGreenLantern and Akaluc expression throughout the parasite life cycle. (A) mGreenLantern expression during asexual development and gametocytes in Pb-mGL-Aka. Thin smears of Pb-mGL-Aka infected erythrocytes were made without fixation, and mGreenLantern fluorescence (green, top panels) was captured using a fluorescence microscope. All mGreenLantern images were captured with the same exposure time. Nuclei were stained with Hoechst 33342 (cyan, middle panels). Phase images are shown below. The bar indicates 10 µm. (B) Representative images of mGreenLantern fluorescence during ookinete maturation. The bar indicates 10 µm. (C) A representative image of mGreenLantern expressing midgut oocysts within a Pb-mGL-Aka infected *A. stephensi* mosquito. A midgut was dissected at day 10 post-feeding and observed using a fluorescence microscope with a phase image. The bar indicates 100 µm. (D) Micrograph of a confocal section of a sporozoite-residing salivary gland. The salivary gland of Pb-mGL-Aka infected mosquito was collected at day 20 post-feeding and stained with Hoechst 33342 to visualize gland nuclei (blue) and CellMask^TM^ Orange (red) to visualize plasma membranes. Fluorescent images are shown as max-intensity projections of confocal stacks. The bar indicates 100 µm. (E) Representative images of an mGreenLantern expressing sporozoite. Sporozoites collected in Figure 7C were stained with Hoechst 33342 (cyan) and observed using a fluorescence microscope. The bar indicates 10 µm. (F) The comparison of oocyst number on mosquito midguts between PbANKA and Pb-mGL-Aka. Midguts were collected at day 10 post-feeding and stained with mercurochrome to count. Each dot represents the number of oocysts from an individual midgut with horizontal lines indicating the median. The Mann-Whitney test was used to compare distributions (N = 30). (G) The number of Pb-mGL-Aka sporozoites collected from mosquito salivary glands were comparable to those of the PbANKA parental line. Fifteen pairs of salivary glands were collected at day 20 post-feeding from PbANKA or Pb-mGL-Aka infected mosquitoes, and sporozoites were counted that successfully invaded salivary glands. Each dot represents the number of pooled sporozoites from five mosquitos. Bars show the mean numbers from biological triplicates with error bars indicating standard deviations. The Mann-Whitney test was used to compare distributions (N = 3). (H) 1×10^4^ sporozoites of the Pb transgenic parasite were injected into an ICR mouse and liver stage development was visualized by measuring luciferase activity 24 hours after inoculation. (I) Comparison of parasite growth in infected C57BL/6 mice between PbANKA and Pb-mGL-Aka parasites. A total of 2000 PbANKA or Pb-mGL-Aka sporozoites were intravenously injected into C57BL/6 mice, and blood parasitemias were determined using Giemsa-stained blood smears. Parasitemia levels of Pb-mGL-Aka parasites (green) were comparable to those of the parental PbANKA strain (black). Error bars represent the standard error of the mean (SEM) from five mice.

## DISCUSSION

In this study, we introduced an mCherry expression cassette into an intergenic locus of the *P. falciparum* NF54 line using the CRISPR/Cas9 system. The resulting NF54-mCh line exhibits robust fluorescence throughout the parasite life cycle, encompassing the blood, mosquito, and liver stages. The strong fluorescence of oocysts facilitates the assessment of parasite transmission to mosquitoes, which enables simpler quantification of the efficacy of transmission-blocking antibodies. The intense fluorescence of salivary gland sporozoites allows for visual detection of infected mosquitoes without dissection, thereby aiding mosquito dissection procedures. In the liver stage, NF54-mCh exhibits robust and constitutive fluorescence from the early stages of infection, with infection rates comparable to the parental line, NF54. The NF54-mCh parasites lacked difficulties observed in other transgenic *P. falciparum* lines, such as insufficient fluorescence in sporozoites and lower infection efficiency in hepatocytes (22,23). To the best of our knowledge, NF54-mCh is the first transgenic parasite line to combine high-intensity fluorescence in salivary gland sporozoites with phenotypically strong development throughout the liver stage. This may be attributed to the *pfhsp70* promoter which was used for driving mCherry expression, and the strategic choice of the intergenic integration locus situated within a transcriptionally active genomic region, which likely ensures stable and high-level transgene expression. The NF54-mCh parasites were able to develop through various life stages *in vitro*, in mosquitoes, and in humanized mice, demonstrating that the introduction of exogenous genes does not interfere with parasite fitness. Because NF54-mCh does not contain a drug resistance cassette, the parasites can be used as a parental line for further genetic modification. In addition, it is notable that no genes were disrupted in the NF54-mCh line because an intergenic locus was used as a target for introducing exogenous genes. For example, NF54-mCh retains an intact *pfs47* locus, which is frequently disrupted in other fluorescent parasite lines; and therefore, its ability to infect refractory mosquito species should be conserved (27). In summary, we believe that NF54-mCh is a valuable tool that will accelerate aspects of *P. falciparum* research, including parasite development and mosquito-parasite interaction.

An exogenous gene expression cassette was also introduced into the orthologous region of the rodent malaria parasites *P. berghei* and *P. yoelii*; and the fluorescence reporter was expressed across all life cycle stages without compromising parasite development. This success indicates that this syntenic locus is a suitable target across different *Plasmodium* species and suggests high versatility regarding the types of exogenous genes that can be introduced. While Kooij et al. previously identified a silent intergenic locus in *P. berghei* as a region suitable for exogenous gene integration (28), the region targeted in this study represents a distinct locus that also supports strong transgene expression. It is likely that additional intergenic loci exist where exogenous gene integration does not impact the parasite life cycle; and utilizing these regions could enable multiple rounds of genetic modification without disrupting any endogenous gene functions. Although we focused on fluorescent protein expression in this study, we believe that these loci could facilitate the introduction of other functional cassettes, such as Cas9 or Cre recombinase; thereby establishing more efficient parental lines for targeted gene editing or inducible gene repression systems. Ultimately, by providing a robust and versatile genetic toolbox, this study offers a transformative platform that will significantly accelerate our understanding of *P. falciparum* biology and facilitate the development of innovative malaria control strategies.

## MATERIALS AND METHODS

### Parasite strains and *in vitro* cultivation

The strain NF54 is known as a high gametocyte producing *P. falciparum* line, and was utilized as the parental strain in this study (39). NF54 parasites were cultivated under 5% CO2 and 5% O2 with human type O erythrocytes at 2% hematocrit in a complete medium consisting of RPMI Medium 1640 (23400-062, Thermo Fisher Scientific, Carlsbad, CA, USA) containing 5.0% human type O serum, 0.5% AlbuMAX II (Thermo Fisher Scientific), 0.225% sodium hydrogen carbonate (FUJIFILM Wako Pure Chemical Corporation, Osaka, Japan), and 0.38 mM hypoxanthine (Sigma-Aldrich, St. Louis, MO, USA) supplemented with 10 µg/ml gentamicin (FUJIFILM Wako). Human serum and erythrocytes were obtained from the Japanese Red Cross Tokyo Blood Center. The parasite was cultured *in vitro* under low oxygen conditions as described (31). For the rodent malaria studies, the *Plasmodium berghei* ANKA and *Plasmodium yoelii* 17XNL strains were propagated in female ICR mice (Japan SLC, Shizuoka, Japan) and used as parental lines for the generation of transgenic parasites.

### Mosquito rearing

In this study, *Anopheles stephensi* SDA500 mosquitoes were used. Larvae were fed a powdered diet (Oriental Yeast Industry, Tokyo, Japan), and adults were maintained on a 3% sucrose solution. Female adult mosquitoes, more than five days after emergence, were allowed to feed on mouse blood and subsequently oviposited. Both larvae and adults were reared at 25°C and 70% humidity under an automatically controlled 16-hour light and 8-hour dark cycle.

### Generation of a transgenic *P. falciparum* line by CRISPR/Cas9-mediated genome editing

Transfection of *P. falciparum* by the CRISPR/Cas9 genome editing system was performed as described (40). The constructions of plasmids for genome editing are described in the Supplemental Materials and Methods. Briefly, to integrate the mCherry expression cassette into the intergenic locus, 25 µg of a centromere plasmid expressing both Cas9 and sgRNA (pCas9/sgRNA) and 25 µg of the linear donor DNA were prepared (see Figure 1A). Synchronized mature schizonts (1.0×10^8^) were purified and mixed with the pCas9/sgRNA plasmid and the donor DNA was dissolved in 100 μl of P3 Primary Cell Solution (Lonza, Basel, Switzerland). Electroporation was performed using the FP-137 program on a 4D-Nucleofector^TM^ device. After electroporation, transfected parasites were cultured in 5 ml of complete medium with fresh erythrocytes, then pyrimethamine drug selection of transgenic parasites was initiated 72 hours after transfection and continued for 10 days. Genome editing of the parasites was confirmed by genotyping PCR using the primer pair pfcheck-F/R. Two clonal parasite lines were isolated by limiting dilution. The nucleotide sequence of the transgenic locus was confirmed by Sanger sequencing.

### Gametocyte culture of *P. falciparum*

Gametocyte culture was performed as described with some modifications (41). Gametocyte cultures were initiated at 0.5% to 0.9% parasitemia with human type O erythrocytes at 5% hematocrit in 5 mL of complete medium per well using 6-well plates. The complete medium consisted of RPMI Medium 1640 containing 50 mg/L of hypoxanthine, 2 g/L sodium bicarbonate (Thermo Fisher Scientific), 2 g/L of D (+)-Glucose (FUJIFILM Wako), and 10% human type AB serum (BioWest, Nuaille, France). To induce gametocytogenesis, parasites were maintained for 15 to 17 days, ensuring that the temperature did not fall below 37°C during daily medium change, but without the addition of fresh human erythrocytes. Gametocytemias and the stages of maturation were determined by Giemsa staining of thin blood smears on glass slides. Male gametocyte maturation was assessed by counting the number of exflagellation centers. To this end, infected red blood cells were adjusted to 10% hematocrit in the serum at room temperature to induce activation and applied to a hemocytometer. After incubation at 20°C for 15 minutes, exflagellation centers were quantified using light microscopy.

### Fluorescence and immunofluorescence microscopy

To visualize fluorescent protein expression in intraerythrocytic stages and activated gametocytes, unfixed thin smears were imaged following nuclear staining with Hoechst 33342. IFAs were performed to identify female and male gametocytes using anti-PfG377 (42) and anti-α-tubulin antibodies, respectively. For female gamete egress assays, anti-Pfs25 and anti-human erythrocyte antibodies were used. To examine liver stage parasites, anti-PfEXP2 (43) was used to label the parasitophorous vacuole membrane. Detailed procedures, including fixation conditions and antibody dilutions, are provided in the Supplemental Materials and Methods.

### Membrane-feeding assay

After 15 to 17 days of gametocyte culture, parasite cultures with robust gametocytemias and exflagellation center assays were fed to mosquitoes using a standard membrane feeding assay as described (40). Briefly, the cultures of *P. falciparum* gametocytes were centrifuged at 900 g for 5 minutes using a centrifuge at 37°C. The supernatant was replaced with human type AB serum, and 50% hematocrit erythrocytes were added to adjust the gametocytemia to 0.5%. Four- to six-day old female *Anopheles stephensi* mosquitoes were fed on gametocytes through Hemotek membrane feeding systems (Hemotek, Blackburn, UK). Female mosquitoes were fed a 10% sucrose solution containing 0.05% gentamicin for 3 days prior to blood feeding, and afterwards the mosquitoes were kept in a 10% sucrose solution at 26°C and 60% humidity. Ookinete and oocyst development were assayed by dissection of mosquito midguts at days 1 and 11 post membrane feeding, respectively. The dissected midguts were ground in RPMI 1649 Medium supplemented with 5 µg/mL Hoechst 33342 to make smears on glass slides for detecting ookinetes. For observation of fluorescent oocysts, dissected midguts were mounted on glass slide and imaged. To assay oocyst numbers, isolated midguts were stained with 0.5% mercurochrome in PBS for 10 minutes, followed by imaging. For imaging of sporozoite-infected salivary glands, the glands were dissected at 18 days post-feeding and stained with 5 µg/mL Hoechst 33342 and CellMask™ Green Plasma Membrane Stain (1:1000 dilution; Themo Fisher Scientific) at room temperature for 5 minutes, followed by imaging with a Leica Stellaris 5 confocal microscope (Leica Microsystems, Wetzlar, Germany). For non-invasive observation of salivary glands with sporozoites, mosquitoes (with legs and wings removed) were anesthetized on ice and imaged using a Leica MZ10 fluorescence stereo microscope. To quantify sporozoites, more than 70 salivary glands were dissected from infected mosquitoes at day 18 post membrane feeding, and homogenized in a 1.5 mL microcentrifuge tube. The number of sporozoites in the homogenates was determined using a hemocytometer under a light microscope.

### Sporozoite infection of human hepatocytes

HC-04 cells (35) were cultured in Dulbecco’s Modified Eagle Medium / Ham’s F-12 (048-29785; FUJIFILM Wako), supplemented with heat-inactivated 10% FBS and 1% penicillin-streptomycin solution (100 U/mL penicillin, 100 μg/mL streptomycin) at 37°C in 5% CO2. The HC-04 cells were transferred to a Nunc Lab-Tek 8-well Chamber Slide (Thermo Fisher Scientific) at 1.0×10^5^ per well and incubated for 24 hours, and then 5.0×10^4^ sporozoites were inoculated into each well. After the addition of sporozoites, wells containing HC-04 cells were cultured in RPMI 1640 Medium (4.5 gm/L glucose, 187-02705; FUJIFILM Wako) supplemented with heat-inactivated 10% FBS, 1% GlutaMAX (Thermo Fisher Scientific), 1% penicillin-streptomycin solution and 1 μg/mL amphotericin B. The culture medium was refreshed every 24 hours.

### Experimental infection of a human liver mouse with sporozoites

A humanized liver TK-NOG chimeric mouse (>70% chimeric) was used for this study and was purchased from CLEA Japan (CLEA Japan, Inc., Tokyo, Japan). A total of 1.0×10⁵ salivary gland sporozoites were suspended in 100 µl of RPMI 1640 medium supplemented with heat-inactivated 2% FBS and were intravenously injected into the chimeric mouse. On days 6 and 8 post sporozoite inoculation, 400 µl of 50% hematocrit human erythrocytes in RPMI 1640 Medium supplemented with 2% FBS was injected intravenously. Four hours after human erythrocyte injection on day 8, blood was collected by cardiac puncture. After thrice washing the collected blood with complete medium, parasites were cultured *in vitro* as described above.

### Rodent malaria parasite growth, mosquito transmission assays, and *in vivo* imaging

Details of procedures to generate transgenic rodent malaria parasites are provided in the Supplemental Materials and Methods. To monitor asexual blood-stage growth of *P. berghei*, C57BL/6 mice (5 to 6 week-old females, Japan SLC) were intravenously inoculated with 2000 sporozoites. For mosquito stage analysis, *A. stephensi* mosquitoes were allowed to feed on infected ICR mice (5 to 6 week-old females) and were subsequently maintained in an incubator at 20°C for *P. berghei* or 24°C for *P. yoelii*. For *P. berghei*, mosquitoes were dissected at 24 hours, 10 days, and 20 days post-blood feeding to examine the fluorescence of ookinetes, oocysts, and sporozoites, respectively. For *P. yoelii*, oocysts and sporozoites were examined 7 and 14 days post-blood feeding, respectively. The salivary glands were dissected and stained at room temperature for 5 minutes with 5 µg/mL Hoechst 33342 and CellMask™ Orange Plasma Membrane Stain (1:1000 dilution; Thermo Fisher Scientific), for 5 min before confocal imaging. For *in vivo* bioluminescence imaging, female ICR mouse were intravenously inoculated with 1.0×10^4^ sporozoites. At 24 h post-inoculation, 100 µL of 5.5 mM Akalumine-HCl (FUJIFILM Wako) was injected intraperitoneally under anesthesia. Images were then taken 10 minutes after the injection using the Newton 7.0 (Vilber, Collégien, France).

## DATA AVAILABILITY

Whole genome sequencing data are publicly available via the DDBJ Sequence Read Archive (DRA) under accession number PRJDB39969.

## ACKNOWLEDGEMENTS

We are grateful to the Japanese Red Cross Society for providing human erythrocytes and plasma. We thank Prof. Sattabongkot (Mahidol University) for HC-04 cells; Dr. Long (NIH) for *P. falciparum* NF54; and Dr. Yamamoto (Jichi Medical University) for *A. stephensi* SDA500. We also thank the Department of Parasitology and Tropical Medicine, Institute of Science Tokyo.

This work was supported by JST SPRING, Grant Number JPMJSP2120 (TS); JSPS KAKENHI under Grant number 24KK0149 to TI and 24K02272 to NS; AMED under Grant Number JP24wm0325074 to TI and JP20wm0325018, JP23wm0225036 and JP23wm0325060 to NS.

Author contributions: TS, HI, TI, and NS conceived the study. TS, RK, DK, YO, KI, HA, and NS performed the experiments and analyzed the data. TS, NS, and TI wrote the manuscript. All authors reviewed the article and approved the final manuscript.

## ETHICS APPROVAL

Ethical approvals for the use of human erythrocytes and plasma from the Japanese Red Cross were granted in accordance with the Medical Research Ethical Committee of the Institute of Science Tokyo. All animal experiments were approved by the Institutional Animal Care and Use Committee of the Institute of Science Tokyo.

